# The Oscillatory ReConstruction Algorithm (ORCA) adaptively identifies frequency bands to improve spectral decomposition in human and rodent neural recordings

**DOI:** 10.1101/855288

**Authors:** Andrew J Watrous, Robert Buchanan

## Abstract

Neural oscillations are routinely analyzed using methods that measure activity in canonical frequency bands (e.g. alpha, 8-12 Hz), though the frequency of neural signals is not fixed and varies within and across individuals based on numerous factors including neuroanatomy, behavioral demands, and species. Further, band-limited activity is an often assumed, typically unmeasured model of neural activity and band definitions vary considerably across studies. These factors together mask individual differences and can lead to noisy spectral estimates and interpretational problems when linking electrophysiology to behavior. We developed the Oscillatory ReConstruction Algorithm (“ORCA”), an unsupervised method to measure the spectral characteristics of neural signals in adaptively identified bands which incorporates two new methods for frequency band identification. ORCA uses the instantaneous power, phase, and frequency of activity in each band to reconstruct the signal and directly quantify spectral decomposition performance using each of four different models. To reduce researcher bias, ORCA provides spectral estimates derived from the best model and requires minimal hyperparameterization. Analyzing human scalp EEG data during eyes open and eyes-closed “resting” conditions, we first identify variability in the frequency content of neural signals across subjects and electrodes. We demonstrate that ORCA significantly improves spectral decomposition compared to conventional methods and captures the well-known increase in low-frequency activity during eyes closure in electrode- and subject-specific frequency bands. We further illustrate the utility of our method in rodent CA1 recordings. ORCA is a novel analytic tool that will allow researchers to investigate how non-stationary neural oscillations vary across behaviors, brain regions, individuals, and species.

## Introduction

Neural oscillations are increasingly recognized as important mesoscopic components of the neural code (Buzsaki 2012; Hanslmayr et al., 2012; Watrous et al., 2015a). Several lines of evidence across species and behaviors demonstrate that the frequency of neural oscillations varies across individuals and shifts to support neural communication and influence behavior (Klimesch 1999; Rudrauf et al., 2006; Cohen 2014; Wutz et al., 2018; Watrous et al., 2013; Furman et al., 2018; Mireau et al., 2017). Across-study differences in both the recording equipment and electrode positioning relative to dipoles may further contribute to frequency variability. Finally, inter- and intra-subject frequency variability has been observed even when using the same equipment and sampling the same cortical areas (Haegens et al., 2014; Zhang et al., 2018). These factors limit researcher’s ability to link oscillations to neuronal spiking and behavior in individual subjects, particularly under circumstances in which frequency variability may obscure spectral decomposition from filtering artifacts (de Cheveigne’ and Nelken, 2019).

To overcome such frequency variability and gain statistical insights by reducing the number of comparisons (i.e. frequencies), many existing approaches perform spectral decomposition in canonical, *a priori* frequency bands (e.g. “alpha”, ~8-12 Hz) and average results over subjects (e.g. Addante, Watrous et al., 2011), although there are several limitations with this approach. First, defining frequency bands can be subject to researcher bias and band definitions are inconsistent across studies, leading to confusion amongst researchers (Newsom and Thiagarajan, 2019). Second, this approach conflates periodic and aperiodic components of the signal and makes assumptions about waveform shape (Haller, Donoghue, Peterson et al., 2018; Cole & Voytek, 2017), and is rarely quantified or compared against alternatives. Finally, the usage of canonical frequency bands obscures subject-level variability.

It thus remains unclear which frequency bands, which we consider as implicit models of oscillatory activity, produce the best spectral decomposition for individual subjects. We posit that the usage of canonical frequency bands has been historically necessary (Brazier et al., 1961) but constitutes an untested model of oscillatory activity that warrants quantification. Building upon prior methods that aim to identify bands based on time-averaged power spectra (Haller, Donoghue, Peterson, et al, Watrous et al., 2018) and quantify oscillatory components of neural signals (Hughes et al., 2012), we sought to derive temporally-resolved and electrode-specific spectral estimates by directly quantifying spectral decomposition performance using different frequency bands.

Here, we present the Oscillatory ReConstruction Algorithm (“ORCA”) that is designed to capture spectral variability and improve spectral decomposition. We introduce new methods for identifying frequency bands based on either spectral peaks relative to the signal background, spectral variability, or explained variance, and compare these methods to canonical frequency bands. ORCA quantifies spectral decomposition performance using each method through signal reconstruction and comparison to the input signal and provides as output the instantaneous amplitude, phase, & frequency of activity in optimized bands. ORCA is thus a novel spectral decomposition and recomposition algorithm that blindly improves spectral estimates using a data-driven approach to minimize experimenter bias. Our results demonstrate that ORCA captures subject- and electrode-specific oscillatory signals in human and rodent data, improves spectral decomposition compared to existing methods, and captures classical low-frequency modulations associated with eye closure in resting scalp EEG. We thus provide a proof of principle for improving the spectral decomposition of diverse neural recordings.

## Results

To investigate the issue of frequency variability across subjects, we first analyzed the frequency content in a scalp EEG dataset recorded from 22 subjects during eyes open and eyes closed resting conditions. We used a reconstruction-based approach that quantifies the explained variance each frequency contributes to the neural signal. Figure 1A shows the r^2^ values for the first 3 subjects in the dataset and reveals considerable diversity in the frequency content of neural signals both across subjects and electrodes. Focusing on occipital sensors across subjects, we nonetheless identified a peak in the canonical alpha range in many subjects and sensors (Figure 1B). Interrogating activity at individual frequencies, we found that average r^2^ values were largest at occipital sites for 10 Hz activity in the canonical alpha band and were largest at frontal midline sites for activity in the canonical delta and theta bands (Figure 1C). Given the considerable frequency diversity across subjects and electrodes (Figure S1), these observations suggest that spectral decomposition should benefit when the particular spectral characteristics of each EEG channel are taken into consideration.

**Figure 1.**
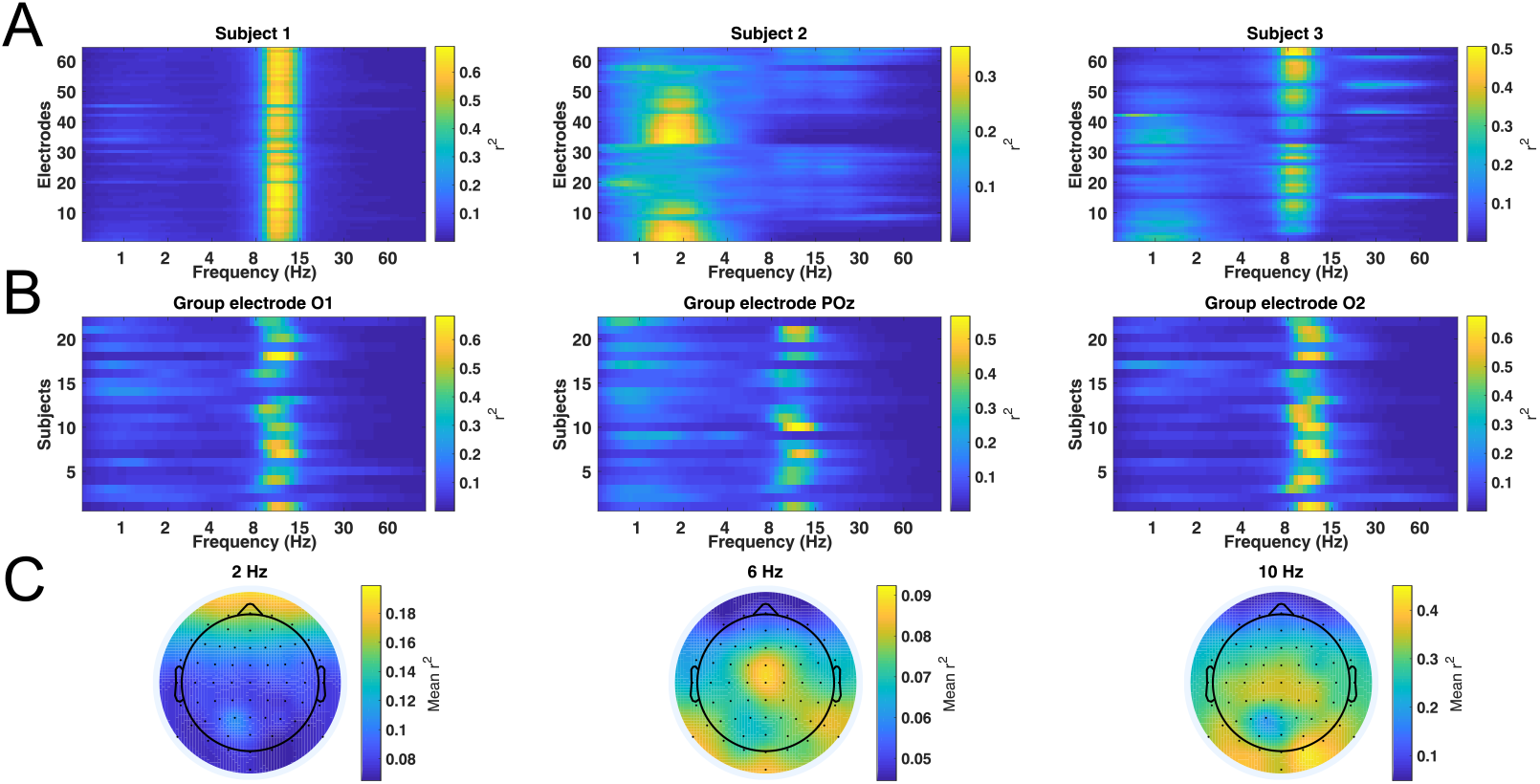
Spectral variability across subjects and electrodes. A) Explained variance (r^2^) at each electrode and frequency in the first 3 subjects. B) Explained variance for all subjects at 3 posterior electrodes, 01, POz, and 02. Most subjects show a peak in explained variance around 10 Hz but with considerable frequency variability across subjects. C) Group-averaged r^2^ values at each electrode location, plotted separately for activity at 2, 6, and 10 Hz. Average r^2^ values are largest over frontal sites at 2Hz and over posterior sites at 10 Hz.

We developed ORCA towards this goal, aiming to improve spectral decomposition by using data-driven band identification methods. Figure 2 shows a schematic of the keys steps in the ORCA algorithm for electrode 01 from subject 1 (see methods for further details). The signal is pre-processed and subject to four different methods for band identification (Figure 2A-B). ORCA uses a subset of the recorded signal to identify bands and avoid over-fitting. The signal is band-pass filtered in each band (Figure 2C) and the amplitude, phase, and frequency of the signal in each band are extracted following a Hilbert transform. These spectral estimates are then used during *spectral recomposition* to reconstruct the input signal (Figure 2D). Reconstruction accuracy is quantified via r^2^ fit between the input and reconstructed signal (Figure 2E). The bands and spectral estimates that produce the best reconstruction are retained and used to calculate a normalized amplitude measure in each band (Figure 2F). On this example electrode, bands based on the explained variance (i.e. Coeffecient of determination method, ‘CoD’, green) outperformed each other method in reconstructing the neural signal (Figure 2E). This example, along with another using rodent data (Figure 2, Supplement 1; see further below for rodent results), quantitatively demonstrates that spectral decomposition can be improved using electrode-specific frequency bands.

**Figure 2.**
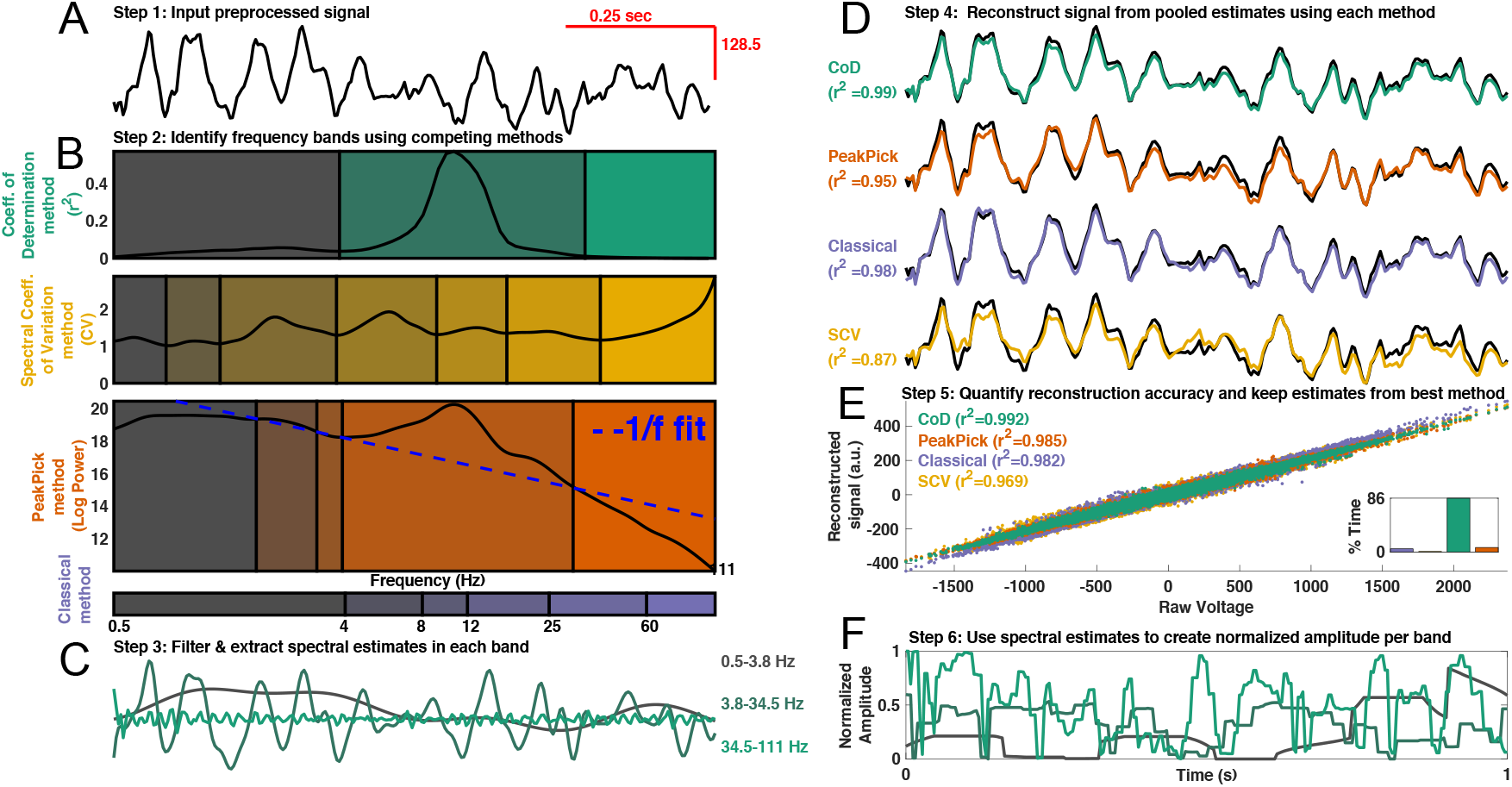
ORCA Schematic. Schematic of the key steps in the ORCA algorithm, illustrated using example electrode 01 from Patient 1. For an example using rodent data, see Figure 2 Supplement 1. A) Step 1: Preprocess the signal, with optional steps including signal rectification, notch filtering to remove line noise, broadband filtering to restrict signal activity into a range of interest (e.g. .5-128 Hz), and downsampling. B) Step 2: Band Identification. Colored boxes indicate different possible band boundaries identified using each of 4 different methods. The upper green panel shows bands identified using the Coefficient of determination (‘CoD”) method, which proved best on this electrode. The middle panel in yellow shows band identified using the spectral coefficient of variation and bands detected using a peak-picking algorithm are shown in blue. The orange panel shows bands identified using the PeakPick method (Watrous et al., 2018). The dashed blue line shows the estimated 1/f signal. The purple lower panel shows the classical bands. C) Step 3: For each method, spectral estimates are extracted in each band using the filter & Hilbert transform method. The filtered signal in each CoD identified band is shown. D) Step 4: Signal reconstruction using spectral estimates derived from each band identification method. Colored traces show the reconstructed signal using each band identification method. The black trace from panel A is superimposed for comparison. R^2^ values indicate the explained variance of the reconstructed data segment to the input data segment. E) Step 5: Reconstruction quantification. Scatter plot shows the raw vs. reconstructed signal for the entire recording for each reconstruction method. Inset bar graph shows the proportion of time each band identification method had the largest r^2^. F) Step 6: Filtered signals and spectral estimates from the method with the highest reconstruction accuracy are used to compute normalized amplitude in each band. The plot shows the normalized amplitude in each band defined using the CoD method.

We ran ORCA on each electrode, first asking which band detection method yields the highest reconstruction accuracy. Figure 3A shows the best method for each subject and electrode and reveals that custom frequency bands outperform classical frequency bands in 93% of electrodes. More specifically, we found that bands defined using the CoD method were best across 57.8% of electrodes, followed by PeakPick in 30.7% of electrodes. The spectral coefficient of variation (SCV) method was best in 4.4% of electrodes and the classical bands were best in 6.8% of electrodes. The CoD and PeakPick methods were best over frontal and posterior channels, respectively (Figure 3, Supplement 1). Assessing electrodes for which each method was best, the CoD, PeakPick, and SCV methods identified an average of 4.1, 4.03, and 4.3 frequency as compared to the classical 6 frequency bands. This observation rules out the possibility that the data-driven methods were superior because they used more parameters (i.e. frequency bands) to reconstruct the signal. Together, these findings indicate that data-driven methods to identify frequency bands can improve spectral decomposition and argue against the usage of *a priori* frequency bands when performing spectral decomposition.

**Figure 3:**
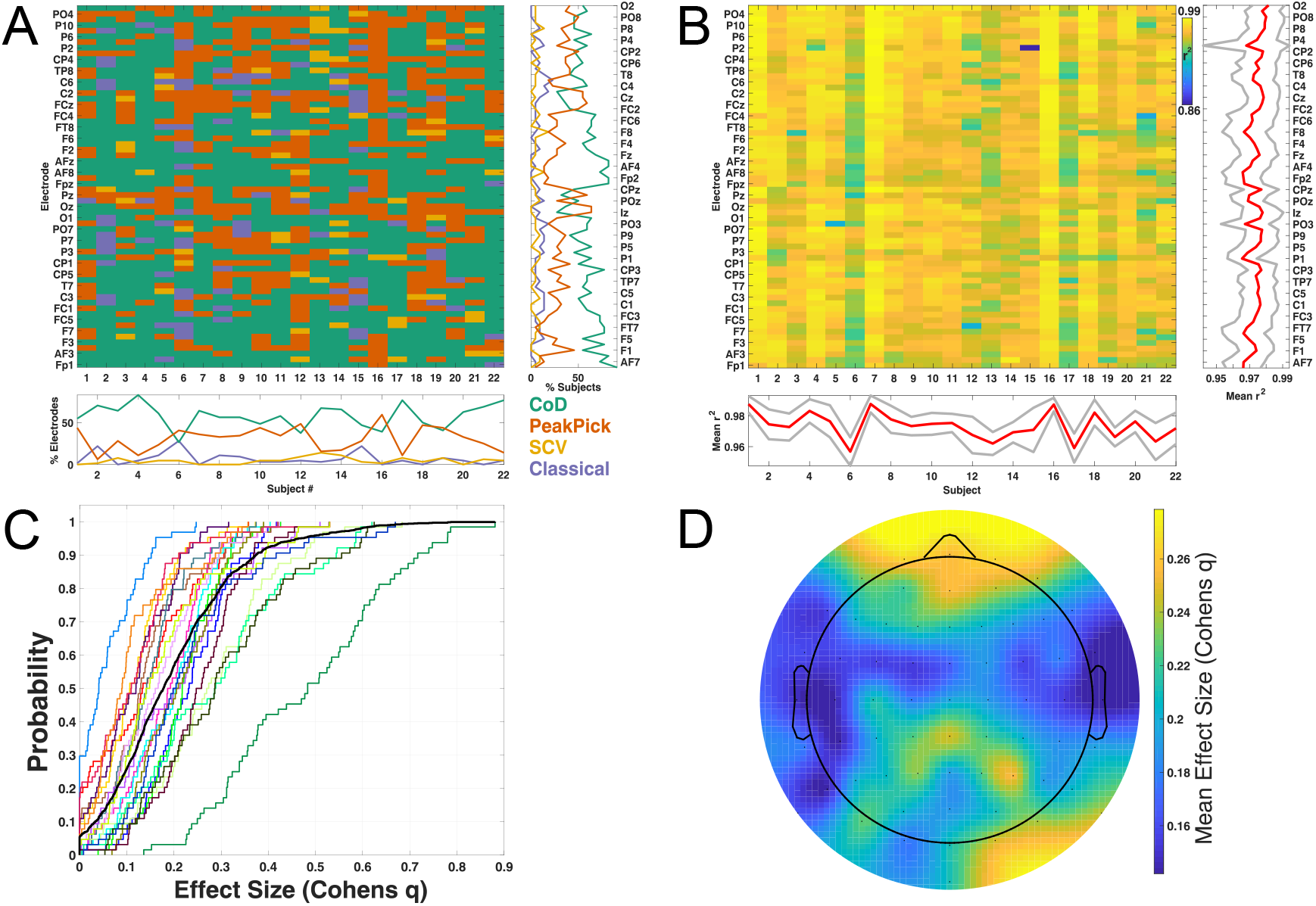
ORCA improves spectral decomposition. A) The winning band-identification method that yields the highest reconstruction accuracy for every subject and electrode. The COD method produces the best results overall, both quantified across subjects at each electrode (right panel) or across electrodes within subject (lower panel). B) Reconstruction accuracy for the winning method for every subject and electrode. The right and lower panels show the mean (red) ± 1 standard deviation (gray) for each electrode and subject. C) Comparison of the best method versus classical frequency bands, expressed as an effect size. Curves show cumulative probability density functions of effect size for each subject. The black line indicates data pooled over all subjects and electrodes. D) Scalp plot showing the average effect size (Cohen’s q) across subjects at each scalp location.

We next investigated the improved performance of ORCA, which was able to capture 97.3% of the signal variance on average when using the best method on each electrode (Figure 3B). We then quantified the improvement in spectral decomposition between different methods. After Fisher’s z-transform, we compared r^2^ values from the best method vs. classical bands (Figure 3B), and observed significantly greater r^2^ values for the best method (paired t-test, t(1407) = 52.7, p<10^-10). Similarly comparing the effect size of improvement between the best and classical band methods on each electrode, the majority of electrodes (74%) showed a small to medium effect size, with substantial variation across subjects (Figure 3C). All but subject 6 showed at least one electrode with a medium effect size (q>.3) and 9/22 subjects showed at least one electrode with a large effect size (q>.5). Frontal and occipital channels showed the largest improvement (Figure 3D). These results demonstrate that ORCA is a superior alternative to conventional methods for spectral decomposition of neural data.

Thus far, we have shown that ORCA improves spectral decomposition through the identification of electrode-specific frequency bands. We next determined if it is feasible to make group level inference using these customized frequency bands on each channel by asking how activity was modulated during eyes open and eyes closed conditions. Figure 4A shows an example electrode whose ~10 Hz activity was significantly modulated during eye closure (Bonferroni p<.05 following permutation test). ORCA captured similar activity modulations spanning the classical alpha and beta bands at most posterior electrodes in this subject (Figure 4B).

**Figure 4.**
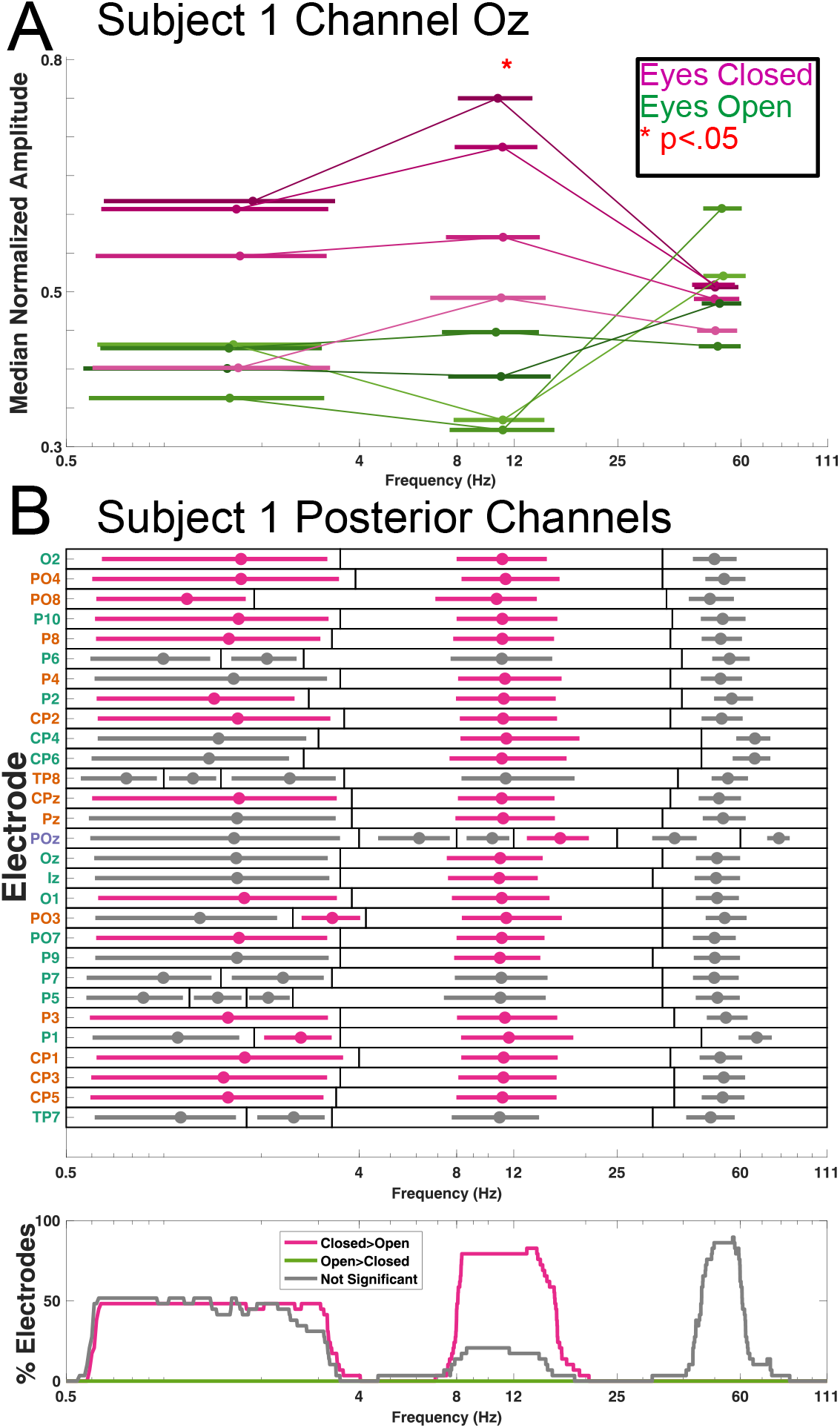
ORCA captures subject and electrode-specific activity modulations during eyes-open and eyes closed conditions. A) Example electrode (Subject 1, electrode 0z) which showed increased amplitude ~10 Hz oscillations during the eyes-closed condition. Thick lines indicate 95^th^ percentile confidence interval for frequencies detected in each band, and each trial is connected with a thin line at the median frequency (small dot). B) Bands detected on each posterior electrode in subjects 1. Horizontal bars indicate 95% confidence intervals for the frequencies detected in each band and are color-coded according to significant differences in the normalized amplitude of activity between eyes open- and eyes-closed task conditions. Black rectangles indicate band edges and dots within each band indicate median frequency of activity. Electrode labels are color-coded by the best band-identification method as in Figure 2 & 3. Lower panel shows the percentage of significant electrodes as a function of frequency.

We observed a similar pattern of results when assessing activity across all subjects at occipital sensors 01 and 02 (Figure 5). Despite heterogeneity in the frequency of activity in these subjects, roughly 80% of subjects showed significant activity increases at 10 Hz during eyes closed conditions over central and posterior electrodes (Figure 5C). These findings indicate that it is possible to understand behavior-related changes in EEG signals at both the individual and group-level using ORCA.

**Figure 5.**
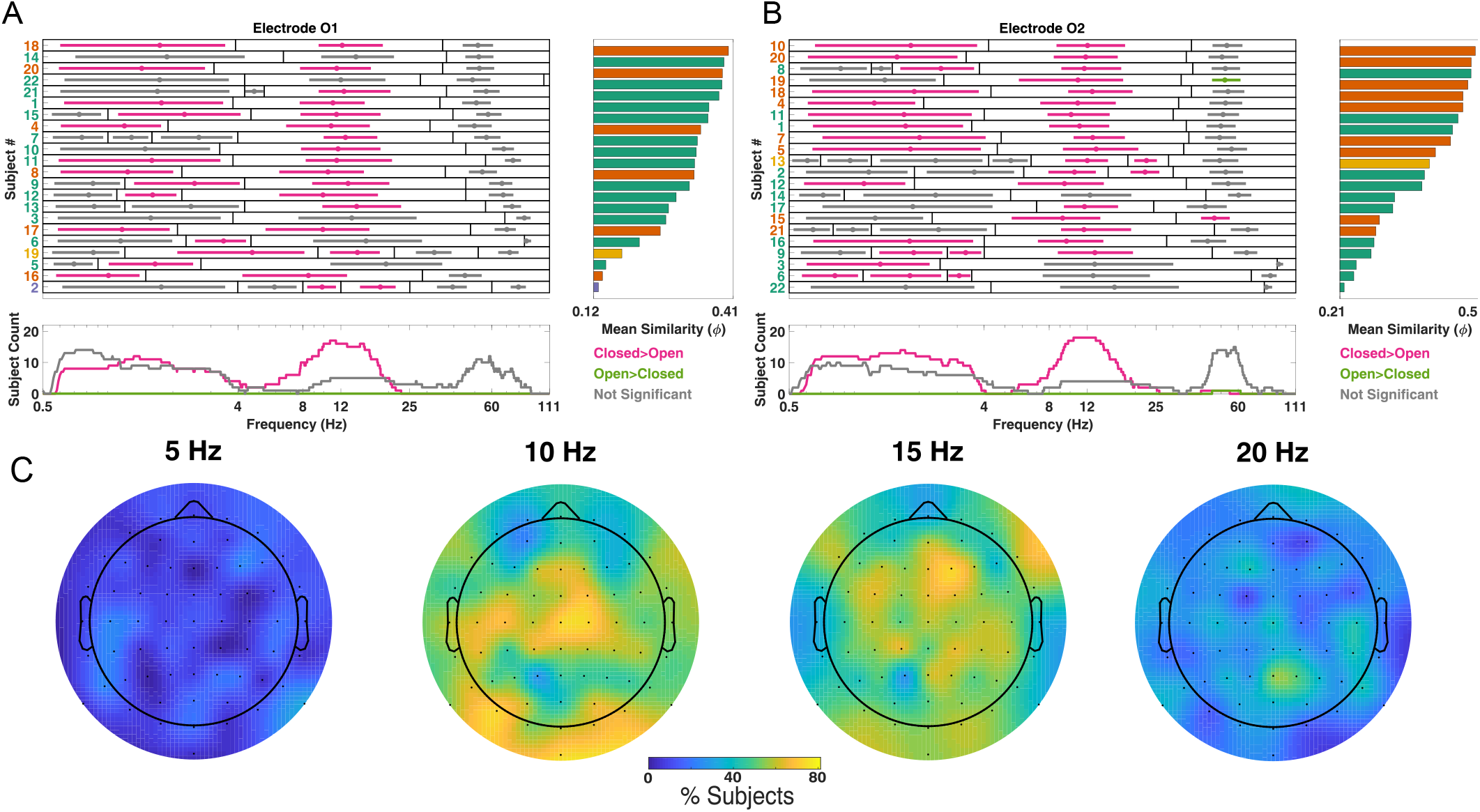
Group level analysis of resting EEG modulations using ORCA. A) Activity modulations in each subject on channel 01. Left panel) Horizontal bars indicate 95% confidence intervals for the frequencies detected in each band and are color-coded according to significant differences in the normalized amplitude of activity between eyes open- and eyes-closed task conditions. Black rectangles indicate band edges and dots within each band indicate median frequency of activity. Electrode labels are color-coded by the best band-identification method as in Figure 2 & 3 and subjects are sorted according to average similarity of their activity to other subjects (right panel; see also Figure 5 Supplement 1). Lower panel shows the percentage of significant subjects as a function of frequency. B) Similar to A, but depicting bands for all subjects recorded at electrode 02. C) Scalp headmaps showing the percentage of subjects showing significant modulation of activity at each frequency. Most subjects showed modulation at 10 Hz over posterior electrode sites.

We next sought to quantify individual differences in the frequency content of neural activity using the output of ORCA. We calculated the inter-subject correlation between the frequency of detected activity in each subject (Figure 5 A-B right panels; Figure 5 Supplement 1). This analysis revealed that the most prototypical subject (Subjects 18 and 10 in Figure 5A and B, respectively) showed activity with median frequency centered slightly above 12 Hz that would likely go undetected using a fixed definition of “alpha activity”. These results further highlight the utility of ORCA in revealing individual differences in neural signals.

We asked how well ORCA performs using other types of neural recordings and analyzed data from rodent hippocampal area CA1 (PFC-2 dataset, crcns.org, Fujisawa et al., 2008). We observed similar performance as in our human dataset (Figure 6), finding that signals on most channels were best reconstructed using the CoD method rather than canonical frequency bands (Figure 2, Supplement 1). ORCA adaptively identified activity in the canonical “theta”, “slow gamma”, and “fast gamma” ranges (Colgin 2016) on most channels (Figure 6B). These results suggest that ORCA can be used on many types of neural signals that are recorded at different spatial scales.

**Figure 6:**
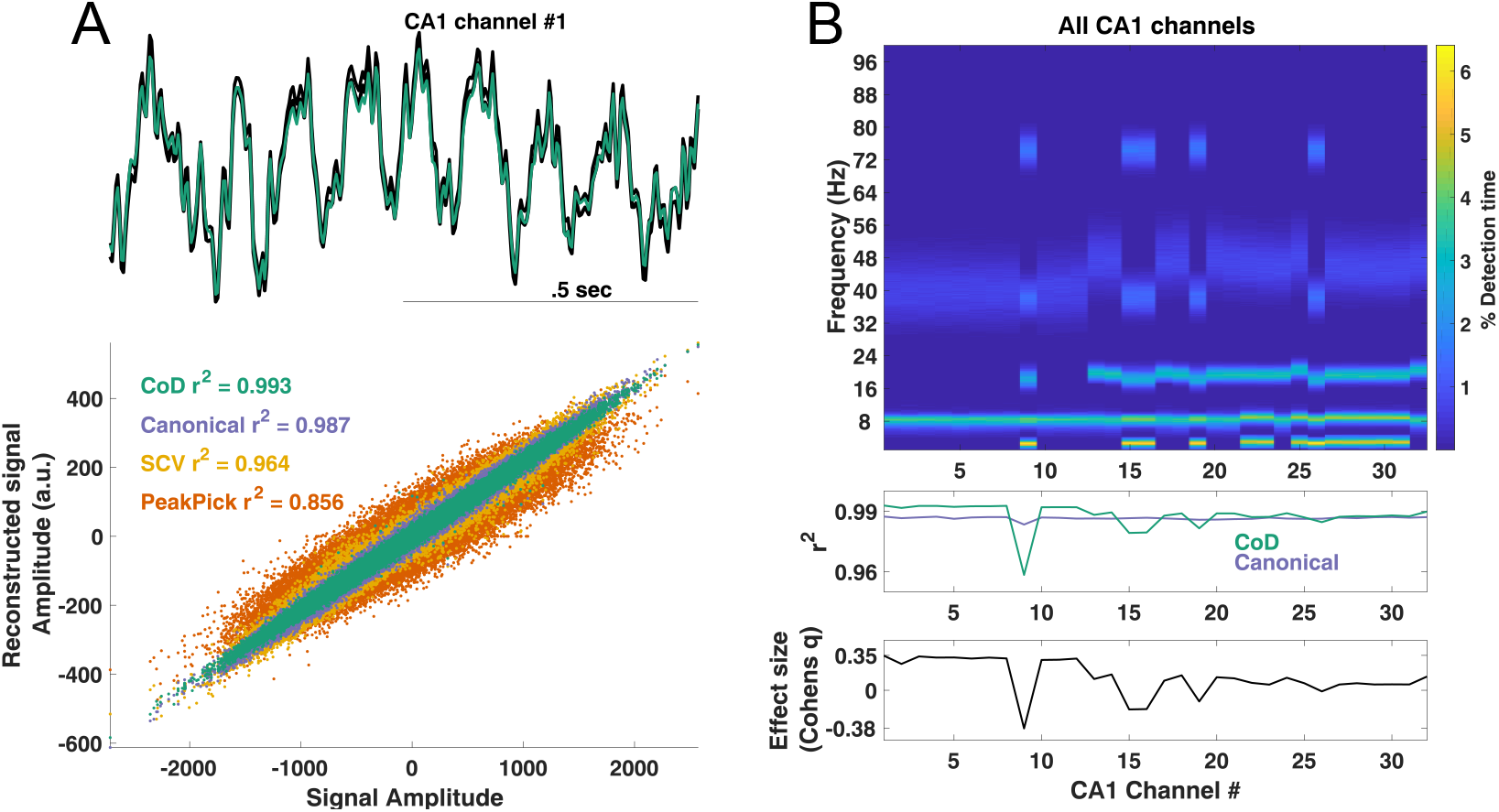
Analysis of rodent CA1 recordings using ORCA. A) Upper: Example raw signal from channel 1 (black) along with the reconstructed signal using CoD bands (green). Lower: Scatter plot showing the raw signal against the reconstructed signal using each band identification method. B) Upper: Detection time as a function of channel number (x-axis) and frequency (y-axis). Middle: r^2^ values for each channel for the CoD and canonical band methods. Lower) Effect size of the reconstruction improvement for the CoD method relative to using canonical bands.

Finally, we performed several control analyses. We quantified the view that band boundaries should be placed far from the signal of interest (de Cheveigne’ and Nelken, 2019), finding that r^2^ values are diminished when a band boundary is located at the same frequency which explain the most signal variance (Figure 6, Supplement 2). We performed a split-halves analysis in the scalp EEG data and found a strong positive correlation (r=.72) between r^2^ values derived separately on the first half and second half of each recording, indicating that oscillatory bands are mostly stable. Lastly, we shifted amplitude and phase estimates in time prior to signal reconstruction in order to test the temporal precision of ORCA and the validity of its output, finding that signal reconstruction is greatly reduced under these circumstances (Figure 6, Supplement 3). Taken together, our results demonstrate that ORCA provides improved spectral estimates in both time and frequency in both human and rodent data, providing a proof-of-principle for future work assessing how electrode and band-specific oscillatory activity co-varies with behavior in spectrally-diverse neural signals recorded in different scales and species.

## Discussion

Analyzing resting EEG and rodent hippocampal recordings, we demonstrate substantial spectral variability across electrodes and subjects in a comparatively simple behavioral setting, highlighting the need for refined approaches when analyzing oscillations. To this end, we developed several novel methods for identifying frequency bands based on different statistical properties of each recording. These methods are incorporated into ORCA, a novel algorithm that pits different models of oscillatory activity against one another to best capture spectral variability and provide improved spectral decomposition. Notably, 93% of channels showed improved spectral decomposition using these new methods rather than canonical frequency bands (Figure 3). ORCA readily identified amplitude modulations in electrode-specific frequency bands associated with eye closure, consistent with decades of research (Berger et al., 1929; Geller et al., 2014; Trujillo et al., 2017). We then applied ORCA to rodent hippocampal recordings and observed that it was capable of blindly identifying theta and gamma components of the neural signal (Colgin, 2016). Our results thus provide a proof-of-principle for using ORCA to analyze electrophysiological recordings with more precision and with less bias than has been previously been possible.

Across all 1408 EEG channels, 93% of channels showed optimized spectral decomposition using customized frequency bands rather than canonical frequency bands. What accounts for such an improvement? We believe this likely occurs because many channels in our EEG dataset have activity spanning the canonical frequency band boundaries (Figure 5, Figure 5 Supplement 1). Given that it is important to select band edges away from the signal of interest in order to avoid filtering artifacts (de Cheveigne’ and Nelken, 2019), it follows that placing a band boundary at 12 Hz using canonical bands would lead to poor filtering and spectral estimation (see also Figure 5, Supplement 1). We conclude that spectral decomposition improvements rendered by ORCA are dependent on the spectral content of the underlying data and thus other datasets may not see such a dramatic improvement in spectral decomposition. Nonetheless, our results clearly argue against the use of canonical, “one-size-fits-all” frequency bands when performing spectral decomposition, and provide a benchmark for quantifying different oscillatory models and spectral decomposition performance through signal reconstruction.

We use the term “optimized” to refer to relative increases in reconstruction between different frequency band models. Going forward, band detection is modular such that improved methods for detecting bands may be incorporated, as was the case with the “CoD” and “SCV” band identification methods. Future work might use genetic algorithms to further refine band identification and/or more explicitly model aperiodic components of the signal (Haller, Donoghue, Peterson et al., *BiorXiv*). We observed that frequency bands were mostly stable over time, though this warrants further investigation, particularly in datasets that contain diverse behavioral states. Future work may extend ORCA by defining bands on smaller subsets of data which could be useful for data cleaning such that data segments with poor decomposition, perhaps based on non-physiological artifacts, can be excluded based on statistically principled grounds.

ORCA draws inspiration from and builds upon prior work which aims to identify and quantify oscillatory components of neural signals, such as “Pepisode”/ “BOSC”, “fooof”, “bicycle”, and “MODAL” (Hughes et al., 2012; Haller, Donoghue, Peterson, et al 2018; Watrous et al., 2018; Cole & Voytek, 2019). Our approach blindly identifies oscillatory bands with very minimal parameterization, quantifies different models of oscillatory activity to improve spectral decomposition accuracy, and provides time-resolved spectral estimates in each band. These features provide new avenues to standardize analysis procedures across research groups. Comparing activity over subjects, ORCA also allows for the identification of comparatively typical and atypical frequency content of neural recordings that may be useful clinically by providing normative data. Finally, ORCA outputs a relatively low-dimensional, “compact” representation of neural signals by providing time-resolved amplitude, phase, and frequency in each band that can facilitate interrogating the relation between neural spiking, oscillatory activity, and behavior.

## Methods

We first provide a description of the ORCA algorithm before describing its key steps and how it was applied to the example datasets.

### The Oscillatory ReConstruction Algorithm (ORCA)

#### Overview

ORCA was developed in Matlab and additionally requires the wavelet toolbox for signal reconstruction. Matlab code for the algorithm is provided on Github (www.github.com/andrew-j-watrous/ORCA. Figure 2 shows a schematic of the key steps in the ORCA algorithm and we describe optional preprocessing and validation steps further below. ORCA requires an input signal that can be any time-series data, the sampling rate, and a wide-band frequency range to be analyzed (e.g. .5-150 Hz). ORCA segments this broad frequency range into bands using 4 different methods (see below) and the signal is band-pass filtered in each band between the band boundaries (e.g. 3 to 12 Hz). ORCA then calculates spectral estimates (amplitude, phase, and frequency of the filtered signal) in each band. Spectral estimates pooled across bands are then used to reconstruct a signal. To measure spectral decomposition performance, the reconstructed signal is compared to the input signal by calculating the linear fit between signals (“regstats” in Matlab), resulting in an r^2^ value for each band identification method. The band-identification method with the largest r^2^ is considered the “best” method and the spectral estimates and bands from this method are retained, while those from the other methods are discarded.

#### Band identification

ORCA uses up to four methods to determine frequency bands. The first and simplest method allows the user to define frequency bands. In this manuscript, we used this method to investigate spectral decomposition using the classical frequency bands (Figure 2B; purple bar), defined as .5-4 Hz “delta”, 4-8 Hz “theta”, 8-12 Hz “alpha”, 12-25 Hz “beta”, 25-60 Hz “slow gamma” and 60-111 Hz “fast gamma”. Throughout this manuscript, we interchangeably refer to this method as “Classical” and “Canonical”.

Each other method defines band boundaries based on different statistical characteristics of the neural signal. By default, these statistics are computed on the first half of the input signal as a means to cross-validate and avoid over-fitting. The second method, “SCV” (Figure 2B; yellow), uses local minima in the spectral coefficient of variation (SCV), a power-normalized estimate of variability at each frequency. Oscillatory power is calculated using 6-cycle Morlet wavelets at 200 log-spaced frequencies from .5 Hz to the Nyquist frequency. SCV is calculated as the standard deviation of power values divided by the mean over time at each frequency. Band edges are defined as local minima in the SCV function. The rationale for this method is that frequencies with comparatively high variability may contain transient oscillations while frequencies with comparatively low variability can then be taken as band edges. We note, however, that semi-continuous oscillatory signals such as rodent hippocampal theta may violate this assumption.

The “CoD” method (Figure 2B, green bars) calculates the coefficient of determination (r^2^) at each frequency by quantifying the fit between the input signal and a reconstructed signal based on activity at each point frequency. Specifically, following spectral decomposition using a continuous wavelet transform, this method uses the inverse continuous wavelet transform separately at each frequency to reconstruct the input signal and quantifies the fit between the input and reconstructed as above. Band edges are defined as 1) local minima in the CoD function and 2) frequencies in which the explained variance is less than what is expected by chance. The rationale here is to use the CoD function to identify band boundaries as frequencies with comparatively low explained variance to the input signal.

Finally, for comparison to previous approaches (Watrous et al., 2018; Lega et al., 2012; Podvalny et al., 2015), we included a fourth method (“PeakPick”; orange bars in Figure 2B). Using the same power values as in the SCV method, we created a power spectrum by averaging wavelet power values over time and fit a line to this spectrum in log-log space using *robustfit* in Matlab. Frequency band edges were defined as those frequencies in the power spectrum that transitioned above or below this fit. Frequency bands for all methods were constrained to be wider than .5 Hz in order to ensure accurate filtering.

### Filtering

Filtering was performed as in the original “frequency sliding” algorithm (Cohen 2014), with one modification that ensured accurate filtering across a variety of frequency bands with different bandwidths (e.g. .5-1 Hz, .5-50 Hz). We thus optimized the transition bandwidth for each frequency band by filtering using different transition widths (.01-.13, .03 steps) and retained the filtered signal with the largest correlation to the raw signal. This modification was necessary for accurate filtering both very narrow and very wide frequency bands. Similar to previous work, instantaneous frequency estimates arising from phase-slips (Cohen, 2014) that were outside of each frequency band were replaced by NaN (Watrous et al., 2018; eLife).

### Signal Reconstruction and quantification of spectral decomposition performance

Reconstructed signals were generated using a synthetic continuous wavelet transform matrix using the instantaneous amplitude, phase, and frequency of activity in each band and then applying the inverse continuous waveform transform (icwt.m in Matlab). This synthetic matrix is sparse, with only as many non-zero values as detected frequency bands at each time sample, and thus the reconstructed signal amplitude is arbitrarily smaller than the observed signal. Spectral decomposition accuracy was determined by calculating the explained variance (r^2^) between the input and reconstructed signal. We then conducted follow-up analyses investigating the proportion of time each method performed best (e.g. Figure 2E) by calculating r^2^ values in 1 second, non-overlapping windows and identifying the method with the largest r^2^ in each window.

### Normalized amplitude calculation

We calculated a measure of normalized amplitude (Figure 2F) using a cycle-by-cycle approach (Cole & Voytek, 2017). The filtered signal in each band is parsed into half-waves by identifying peaks and troughs in the filtered signal and the amplitude of each half-wave is then calculated as the absolute value of the peak to trough height. To account for the approximately inverse relation between oscillatory frequency and amplitude, we normalized each half-wave amplitude by multiplying it by its instantaneous frequency. Each half-wave is then ranked against all others across the full recording such that all values are within a range of 0 to 1 (smallest to largest, respectively).

### EEG Dataset and analyses

For results related to human recordings, we analyzed a published scalp EEG dataset (EEG; Trujillo et al., 2017) of 22 subjects recorded during eyes open- and eyes-closed conditions. This dataset consists of 64 scalp channels sampled at 256 Hz and referenced to a common mode sense electrode located between sites Po3 and POz. Subject performed a total of 8 minutes of interleaved, 60 second blocks of either eyes-open or eyes-closed conditions (4 “trials” each). For this EEG dataset, we mean-centered each recording and performed line noise reduction using a bandstop filter from 58-62 Hz prior to decomposition with ORCA and did not perform artifact correction.

Each pre-processed channel was analyzed with ORCA as a continuous, unepoched recording. Following spectral decomposition with ORCA, the median normalized amplitude value was extracted from each 60-second trial in each detected frequency band (Figure 4A). These median values for eyes-open and eyes-closed conditions were compared using nonparametric Mann-Whitney tests. We shuffled the condition labels associated with each value a total of 70 times (corresponding to the number of unique groupings of 8 values) and recomputed a pseudo test statistic. The true test statistic was ranked against the distribution of 70 pseudo test statistic values to derive a shuffle-corrected p-value. We then performed Bonferroni correction for multiple comparisons (frequency bands) on each electrode. P-values exceeding the 95^th^ percentile or below the 5^th^ percentile after Bonferroni-correction were considered significant.

To identify subjects with similar activity (Figure 5), we first generated a Boolean matrix for each channel indicating whether activity was detected at each frequency when using different percentile inclusion criteria (Figure 5, Supplement 1). This allowed us to circumvent the issue that each channel may have different numbers of frequency bands and that the same frequency (e.g. 10 Hz) may be included in a different band in different subjects. We calculated the Phi correlation between these Boolean matrices in order to determine similarity of detected activity between subjects. We then calculated the mean Phi coefficient for each subject to determine each subject’s average similarity on each channel.

### Rodent Dataset and analyses

For results related to rodent recordings, we analyzed a subset of recordings from a publicly-available dataset (Fujisawa et al., 2008; crcns.org PFC-2 dataset). Specifically, we analyzed the first 5 minutes of CA1 recordings from session “ee708/EE.188” during which the rat was performing a spatial working memory task. Prior to decomposition with ORCA, signals were downsampled to 312.5 Hz. Signals were broadband filtered from 1-100 Hz and canonical bands were defined as 1-4, 4-12, 12-25, 25-55, and 55-100 Hz (Colgin, 2016). We again implemented a cross-validated band-identification procedure such that the first 2.5 minutes of the signal were used to generate bands for the CoD, SCV, and PeakPick band identification methods. We did not do artifact rejection.

We tested the assumption that band boundaries should be placed far from the signal of interest (de Cheveigne’ and Nelken, 2019) using the first CA1 channel in the rodent recordings. This signal was chosen because it was best reconstructed using two bands and a single frequency boundary at 17.1 Hz (Figure 6, Supplement 1). To this end, we generated a separate model with a single band boundary at each frequency and recalculated signal reconstruction accuracy.

## Supplemental Figures

**Figure 1 Supplement 1.**
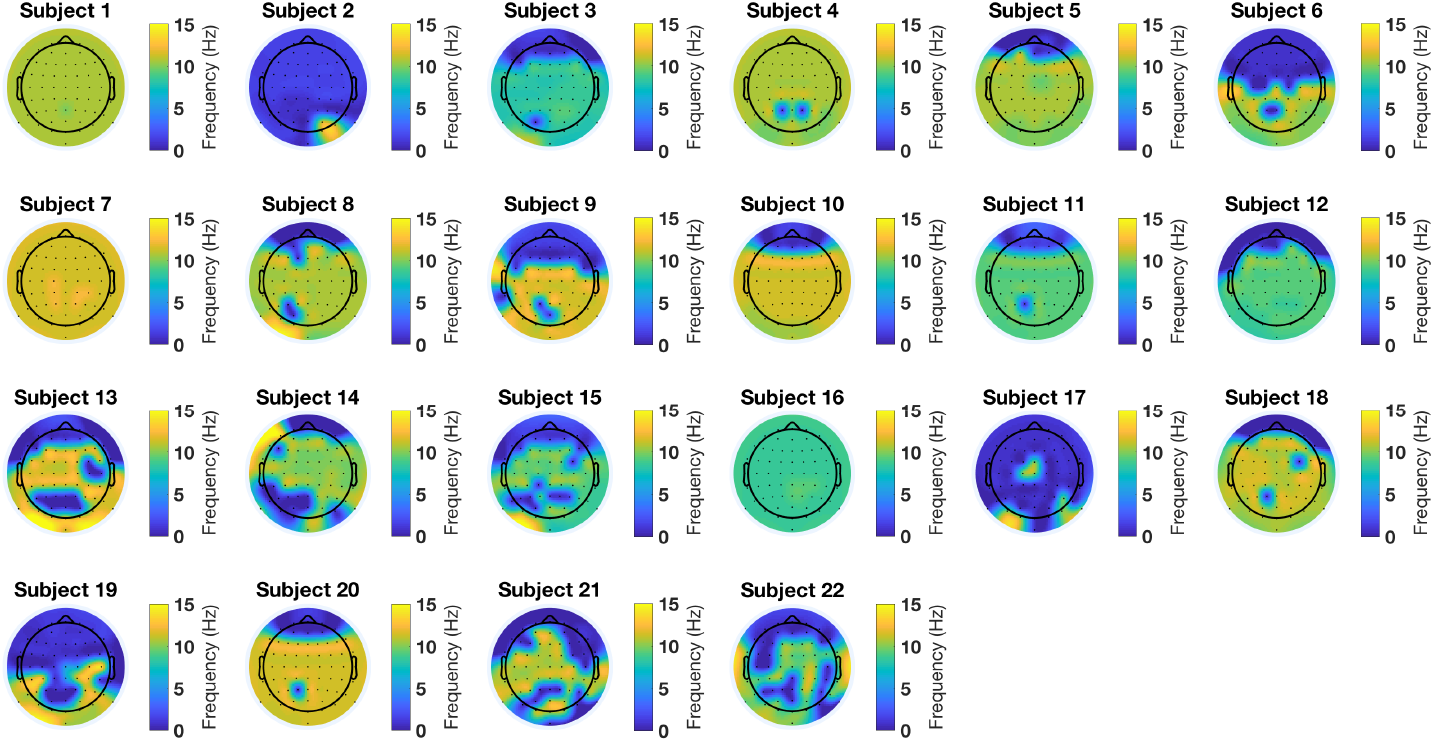
The frequency that explains the most variance to recordings on each electrode is plotted as a scalp topography for each subject. Note that most subjects have different frequencies at different sites and also that some subjects have stable frequencies across locations but differ between themselves (e.g. Subject 1 and 16). These observations motivate the use of spectral decomposition methods that account for frequency variability across individuals and electrodes.

**Figure 2, Supplement 1.**
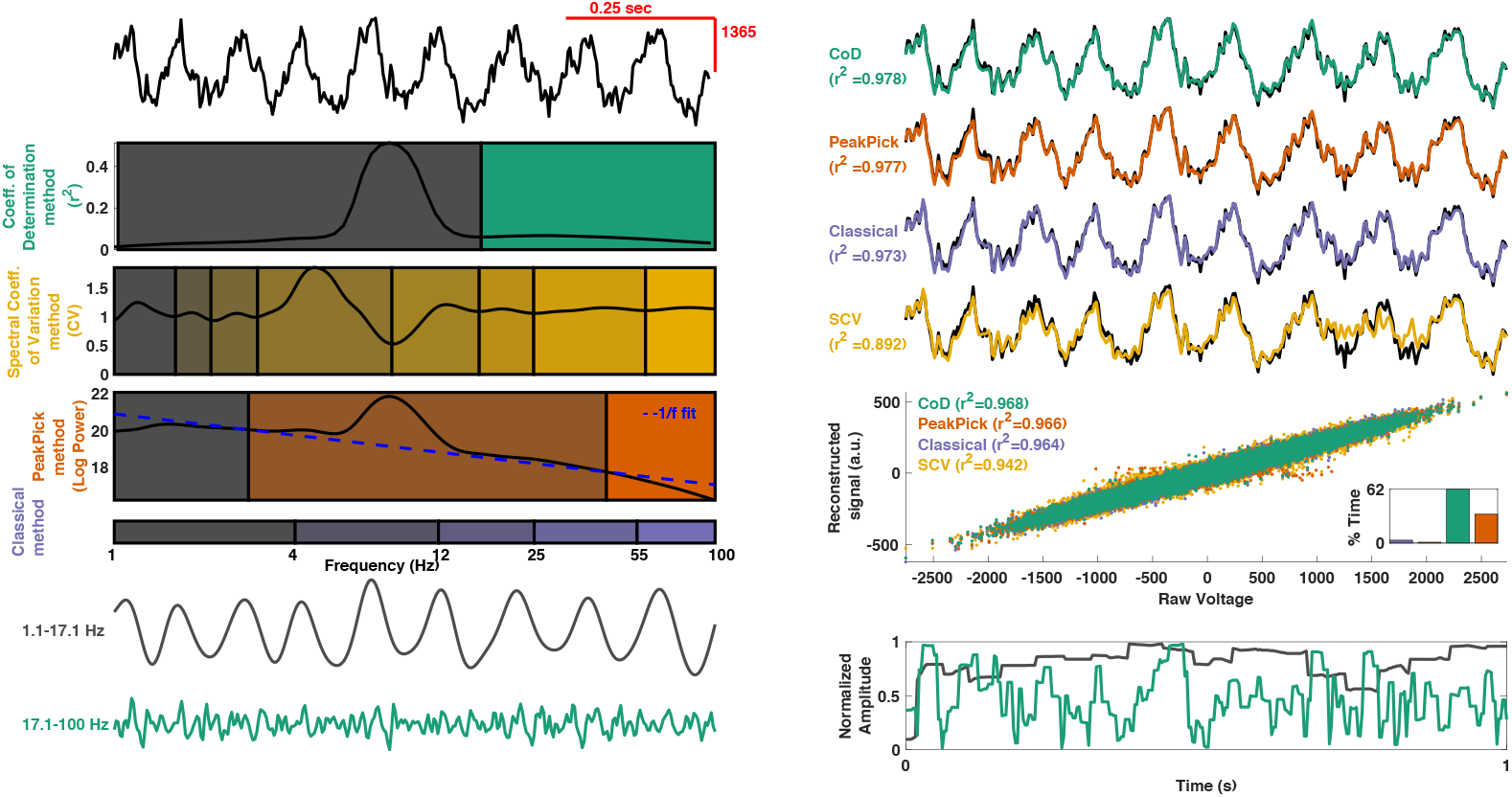
ORCA schematic using channel 1 from the rodent dataset. See Figure 2 caption for further details on figure layout.

**Figure 3, Supplement 1.**
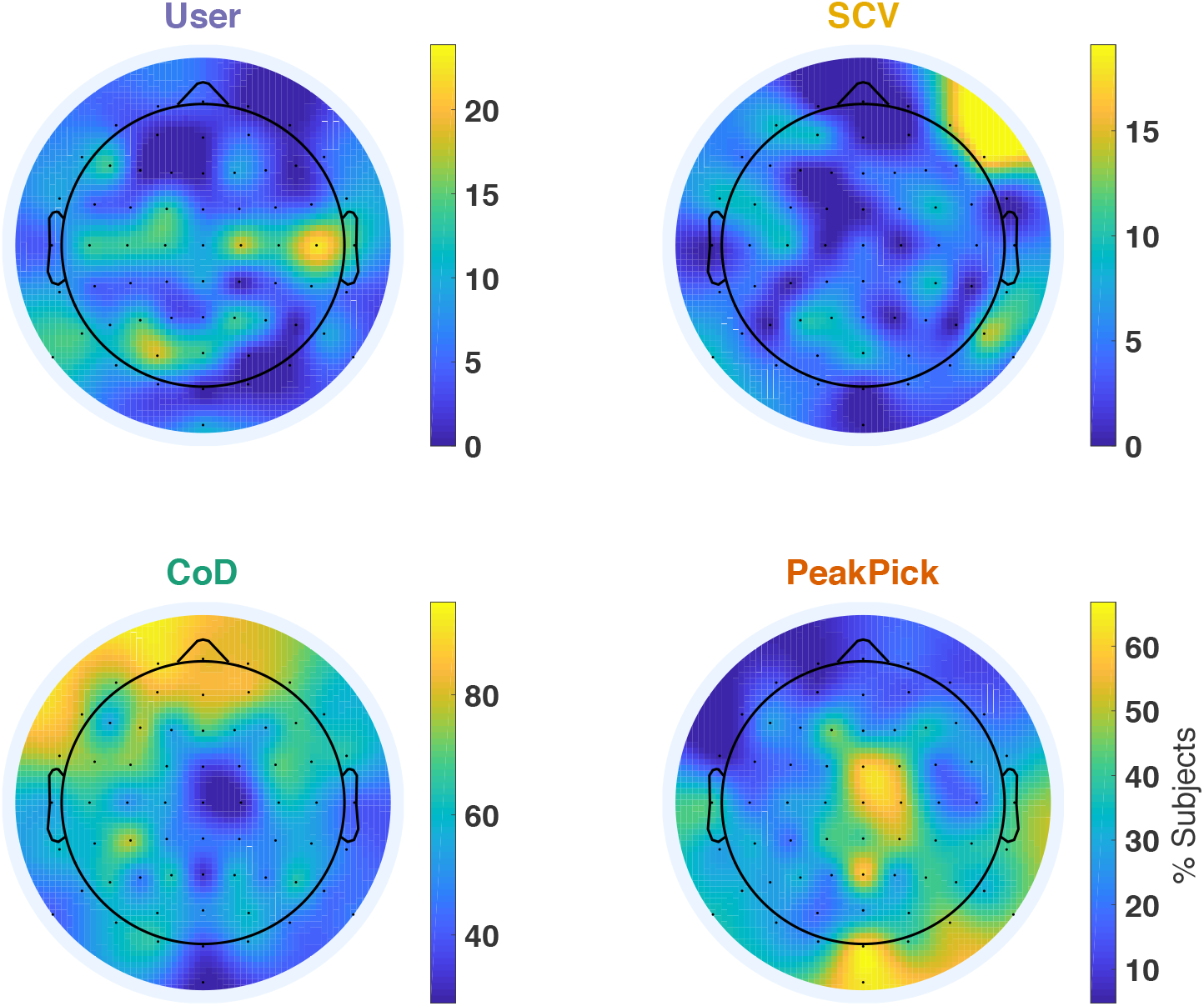
Scalp topographic maps showing the percentage of subjects for which each method yielded the largest reconstruction accuracy.

**Figure 5, Supplement 1.**
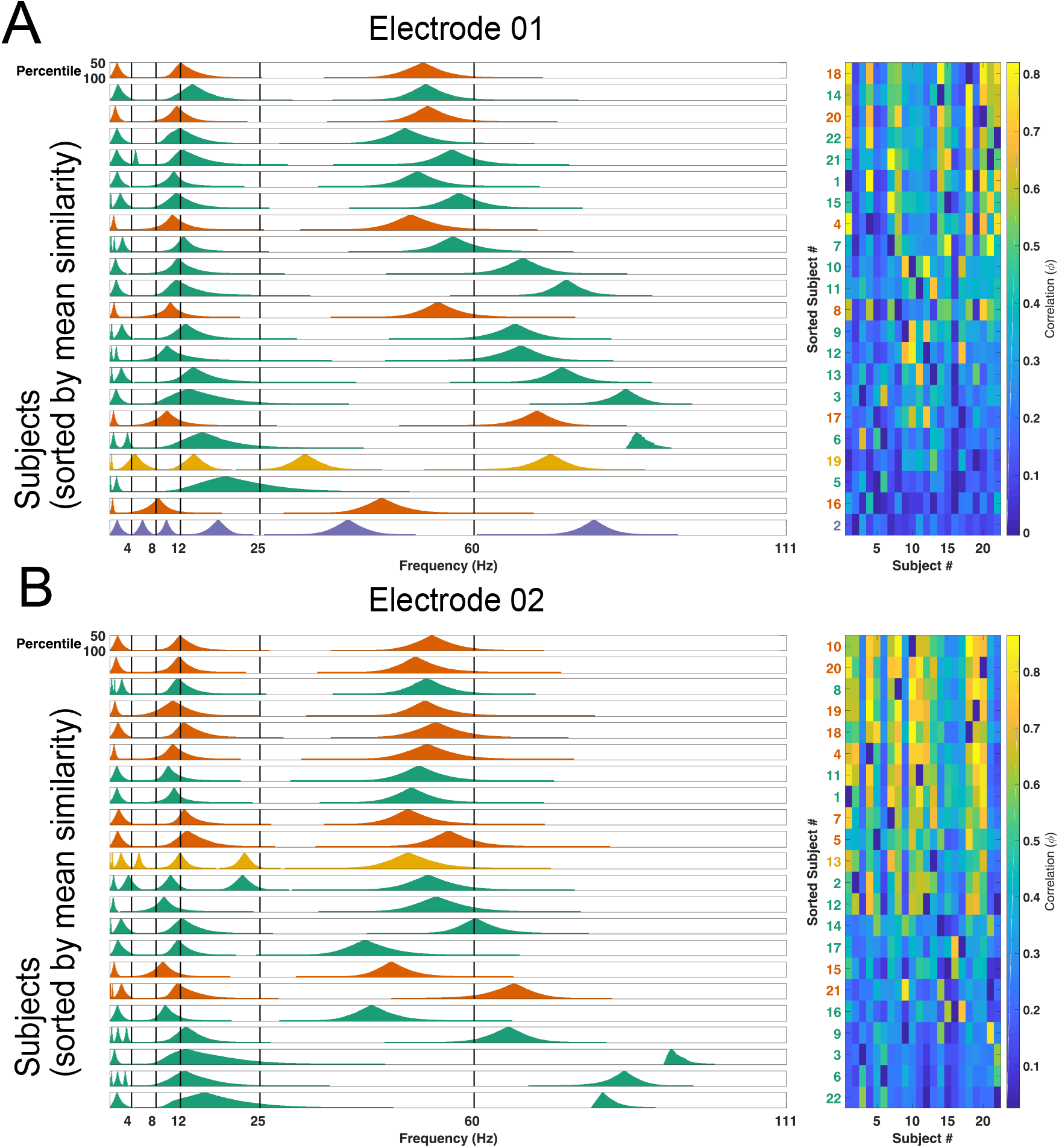
A) Frequency bands for each subject defined over different confidence levels (y-axis in each subpanel) for electrode 01. Data for each subject is color-coded according to the band-identification method used, as in Figures 2 and 3. Note that many subjects have activity spanning the canonical frequency bands (black vertical lines). Right panel shows the inter-subject correlation coefficients for each sorted subject.In both panels, subjects are sorted according to their mean similarity with activity detected in other subjects, from least to most similar (bottom to top, respectively). B) Same layout as A but for electrode 02.

**Figure 6, Supplement 1.**
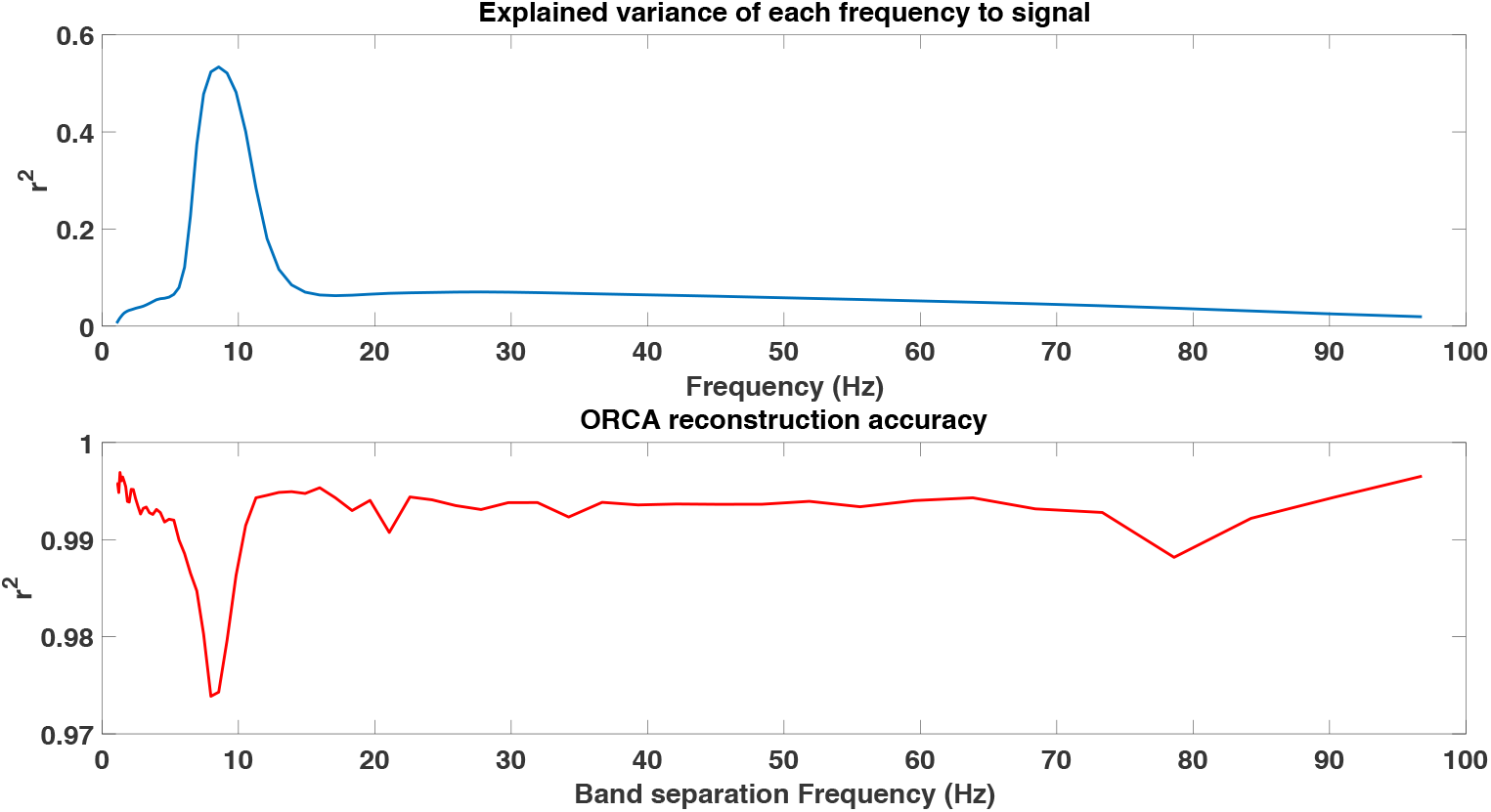
Upper panel: Explained variance of each frequency to the recorded signal for rodent CA1 channel 1. This channel was best reconstructed using a single band boundary at 17.1 Hz (Figure 2, Supplement 1). Lower panel: Reconstruction accuracy as a function of the location of this single band boundary. Reconstruction is worst when the band boundary is placed at ~8 Hz, validating the idea that (filter) band boundaries should be placed away from the signal of interest when performing filtering and spectral decomposition (de Cheveigne’ and Nelken, 2019).

**Figure 6, Supplement 2.**
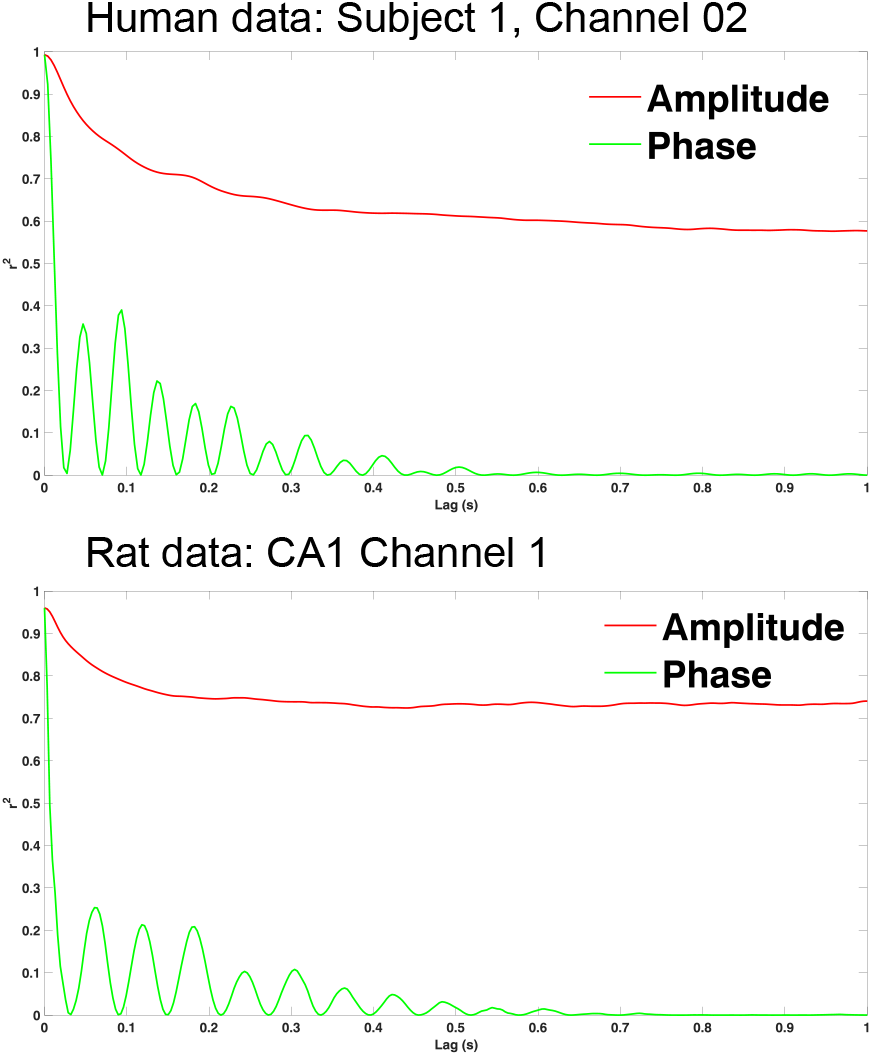
Shifting spectral estimates in time abolishes reconstruction accuracy. Upper) Human data showing that increasing the lag (x-axis) between either the amplitude or phase estimates and the input signal reduces reconstruction accuracy (y-axis). A lag of zero indicates no temporal shuffling for which reconstruction accuracy is > .95. Lower) Same as above but for the first channel of rodent CA1 data.

## Acknowledgments

We thank Arne Ekstrom for helpful feedback on this manuscript.

## Notes

https://github.com/andrew-j-watrous/ORCA

